# Design of multi-epitope vaccine candidate against Brucella type IV secretion system (T4SS)

**DOI:** 10.1101/2023.05.16.540940

**Authors:** Zhengwei Yin, Min Li, Ce Niu, Mingkai Yu, Xinru Xie, Gulishati Haimiti, Wenhong Guo, Juan Shi, Yueyue He, Jianbing Ding, Fengbo Zhang

**Affiliations:** The First Affiliated Hospital of Xinjiang Medical University, No. 393, Xinyi Road, Urumqi 830011, Xinjiang, China; Department of Clinical Laboratory, The First Affiliated Hospital of Xinjiang Medical University, No. 393, Xinyi Road, Urumqi 830011, Xinjiang, China; Department of Immunology, School of Basic Medical Sciences, Xinjiang Medical University,No. 393, Xinyi Road, Urumqi 830011, Xinjiang, China; State Key Laboratory of Pathogenesis, Prevention, Treatment of Central Asian High Incidence Diseases, the First Affiliated Hospital of Xinjiang Medical University, No. 393, Xinyi Road, Urumqi, 830011 Xinjiang, China

**Keywords:** Brucella, Multi-epitope vaccines, T4SS, VirB8, VirB10

## Abstract

Objective the dominant epitopes of T and B cells of VirB8 and VirB10 of Brucella type IV. Secretory systems were predicted and analyzed by bioinformatics, and then multi-epitope vaccines were constructed by reverse vaccination. Methods The amino acid sequences of VirB8 and VirB10 were obtained from the UniProt database, T and B cell dominant epitopes were selected and supplemented with adjuvants to construct a multi-epitope vaccine, SOPMA and RoseTTAFold were applied to predict secondary and tertiary structures respectively for immune simulations. Finally, molecular dynamics simulations of iMEV-TLR4 were then performed. Results one cytotoxic T lymphocyte (CTL) epitope, five helper T lymphocyte (HTL) epitopes, two linear B cell epitopes and three conformational B cell epitopes were obtained in VirB8. 1 cytotoxic T lymphocyte (CTL) epitope, 4 helper T lymphocyte (HTL) epitopes, 4 linear B cell epitopes, and 3 conformational B cell epitopes were obtained in VirB10. The resulting multiepitope vaccine is a protein with good antigenicity, hydrophilicity, and stability. Conclusion a novel brucella multiepitope vaccine was designed by reverse vaccination, and this study provides a theoretical basis for further research.

## Introduction

Brucellae are tiny, globular, gram-negative bacteria that are non-budding, non-flagellated and do not form pods. The genus Brucella includes twelve species: Brucella melitensis, Brucella abortus, Brucella hoggetti, Brucella neoformans, Brucella suis minor bifidum, Brucella catarrhalis, Brucella canis, Brucella vulgaris, Bifidobacterium cetaceum, Bifidobacterium barkerium and Bifidobacterium plumosa^[1]^.Although Bifidobacterium melitensis mainly infects sheep and goats, it often causes brucellosis in humans^[2]^.Brucellosis is one of the most common zoonotic infections worldwide^[3,4]^.The main clinical manifestations are fever, weakness, arthralgia and muscle pain^[5]^.The main routes of transmission are the gastrointestinal tract, skin, mucous membranes, respiratory tract, blood body fluids and aerosols.For example, direct or indirect contact with infected animals, consumption of their raw meat or dairy products^[6,7]^.In recent years, the incidence of human brucellosis in China has increased dramatically^[8]^.Brucellosis can involve multiple organs, but its clinical manifestations are non-specific^[9]^.Traditional bacteriological methods take a long time to identify^[10]^.At present, human brucellosis remains one of the major public health problems in China. For this reason, new treatments against Brucella are being developed in early prevention.

Vaccines are an effective way to prevent brucellosis^[11]^.Reverse vaccinology (RV) has proven to be a very effective approach in which vaccine antigen prediction is performed based on bioinformatics analysis of the pathogen genome using rational vaccine design^[12,13]^.MEV, on the other hand, uses a reverse vaccinology approach to vaccine prediction, which can improve its safety and efficacy^[14]^.The VirB system (VirB1-VirB12) has been shown to be present in all Brucella species and is highly conserved^[15]^.The type IV secretion system (T4SS), encoded by the virB manipulator, is an important virulence factor for Brucella abortus. It disrupts cellular pathways, induces a host immune response by secreting effectors, promotes replication of Brucella in host cells and induces persistent infection^[16]^.VirB8 is one of the core components of the type IV secretion system (T4SS) and has been shown in previous experiments to be immunogenic and suitable as a candidate protein for vaccine design^[17]^.VirB10 is an important functional protein of Brucella abortus and one of the core components of the type IV secretion system (T4SS)^[18]^.VirB10 has been shown to have good immunogenicity in animal models of infection and is a good choice for vaccine design^[19]^.Ultimately, both proteins are suitable for MEV design against Bifidobacterium melitensis.

We analysed the epitopes of VirB8 and VirB10 using various bioinformatics methods such as IEDB, NetCTLPAN1.1, NETMHCIIpan4.0 and ABCpred.Epitope bonding requires a linker. We use AAY, GPPGPG and KK to link CTL epitopes, HTL epitopes and B-cell epitopes^[20]^.However, epitope-linked vaccine peptides are poor immunogens and susceptible to enzymatic degradation, so we chose specific adjuvants to enhance and stabilize the immunogenicity of the vaccine peptides^[21]^.We analyzed the physicochemical properties, antigenicity and sensitisation of MEV, and performed a model assessment of the predicted secondary and tertiary structures of MEV, followed by immune simulations and finally a molecular dynamics study of MEV-TLR4.

## Materials and methods

### 1.1 Material sources

Find the amino acid sequences of Brucella melitensis VirB8 and VirB810 in the UniProt database.

### 1.2 Research Methodology

#### 1.2.1 Selection of target proteins

ProtParam (http://web.expasy.org/protparam/) software was applied to analyse the physicochemical properties of the proteins and MEVs. This included the number of amino acids, the molecular formula, the instability index and the overall mean of the water solubility (GRAVY). VaxiJen (http://www.ddg-pharmfac.net/vaxijen/VaxiJen/VaxiJen.html) was applied to analyse the antigenicity of the target proteins^[22]^with a threshold value of 0.4. For this purpose we also compared hydrophilicity and stability. Ultimately, VirB8 and VirB10, the two target proteins, had good antigenicity, hydrophilicity and stability when designed for MEV.

#### 1.2.2 Sequence Search

The amino acid sequences of VirB8 (sequence number Q9RPX7) and VirB10 (sequence number Q8YDZ0) were obtained from the Uniprot database (https://www.uniprot.org/). The amino acid sequences in Jalview were compared by MAFFT to analyse their homology^[23]^.

#### 1.2.3 Prediction of signal peptides

The prediction of protein signal peptides was implemented by SignalP5.0 and LiPOP1.0(https://services.healthtech.dtu.dk/service.php?LipoP-1.0). finally, we chose the merged set as the final result^[24]^.

#### 1.2.4 Prediction of protein T-cell epitopes

T cell epitopes consist of MHC class I and MHC class II molecules, and CD8 T cells become cytotoxic T lymphocytes (CTL) upon recognition of the T CD8 epitope.At the same time, the triggered CD4 T cells become helper T lymphocytes (HTL) or regulatory (Treg) T cells^[25,26]^.We selected the Xinjiang HF alleles (HLA-A*1101, HLA-A*0201, HLA-A*0301, HLA-DRB1*0701, HLA-DRB1*1501 and HLA-DRB1*0301) to predict CTL, HTL epitopes^[27]^.CTL epitopes of target proteins are predicted by IEDB (http://tools.immuneepitope.org/) and NetCTLpan1 server (https://services.healthtech.dtu.dk/service.php?NetCTLpan-1.1)[28],29].For CTL epitope prediction, the three alleles of HLA-A were selected with a length of 10 and the other original thresholds were unchanged. As NetCTLpan1.1 starts counting from 0, care should be taken to add 1 to the sequence when comparing the results with those of IEDB at the end.Application of IEDB and NetMHC-IIpan-4.0 (https://services.healthtech.dtu.dk/service.php?NetMHCIIpan-4.0) for prediction of HTL epitopes of target proteins^[30]^.For HTL epitope prediction, three allele lengths such as HLA-DRB1 were chosen to be 15.The default thresholds for NetMHC-IIpan-4.0 remain unchanged.Ultimately, we listed the top 10 high-scoring epitopes for each software and selected the two software repeats as T-cell dominant epitopes for the proteins.

#### 1.2.5 Prediction of protein B-cell epitopes

B-cell epitopes consist of linear and conformational epitopes.ABCpred (https://webs.iiitd.edu.in/raghava/abcpred/ABC_submission.html) was used to predict the selection of dominant B-cell linear epitopes with an overall prediction accuracy of 65.93%. The corresponding sensitivity, specificity and positive predictive values were 67.14 %, 64.71 % and 65.61 %, respectively.The prediction of conformational epitopes was performed by Ellipro of IEDB (http://tools.iedb.org/ellipro/), and we selected sequences with high scores. Finally, linear and conformationally dominant epitopes of B cells were selected for MEV design.

#### 1.2.6 Molecular docking of T-cell epitopes to HLA alleles

Use the HDOCK server for molecular docking.We selected HLA class I (HLA-A*02:01) and HLA class II (HLA-DRB1*01:01) alleles for molecular docking with T-cell epitopes and eventually discovered the interactions between the alleles and T-cell epitopes.

#### 1.2.7 Vaccine construction for MEV

We choose non-toxic, non-allergenic and advantageous table positions.CTL epitopes are linked to AAY linkers and linked to HTL epitopes with GPGPG linkers; HTL epitopes are linked to GPGPG linkers and linked to B-cell epitopes with KK linkers; B-cell epitopes are linked with KK linkers ^[20]^. To increase the antigenicity of MEV, the human β-defensin-3 sequence (sequence no. Q5U7J2), and the PADRE sequence were linked with the help of an EAAAK junction at the N terminus^[31]^.Finally, a polyhistidine tag is added to the C-terminus to obtain the complete vaccine protein sequence.

#### 1.2.8 Physicochemical properties, antigenicity, solubility and sensitization of MEV

ProtParam software was applied to analyse the physicochemical properties of MEV. VaxiJen was applied to analyse the antigenicity with a threshold of 0.4 and SOLpro (http://scratch.proteomics.ics.uci.edu/) was applied to predict the solubility of MEV. with a threshold value of 0.5^[32]^. Finally, AllergenFP was applied to analyse its allergenicity.

#### 1.2.9 Projections for secondary and tertiary structures

We used the SOPMA (http://npsa-pbil.ibcp.fr/cgibin/npsa_automat.pl?page=/NPSA/npsa_sopma.html) secondary structure prediction tool. Applying RoseTTAFold to predict the tertiary structure of MEV^[33]^.

#### 1.2.10 Quality assessment of predictive models

The quality of the three-level structural model of the MEV was assessed using the SWISS-MODEL structural assessment (https://swissmodel.expasy.org/assess).

#### 1.2.11 Molecular docking

Molecular docking of MEV and TLR4 (PDB ID: 4G8A) immunoreceptors using the HDOCK server (http://hdock.phys.hust.edu.cn/)[34].Key interacting residues were inferred from LigPlot+ generated protein-ligand 2D structural interaction maps and their 3D structures were visualised by PyMOL.

#### 1.2.12 Immunosimulation

C-ImmSim (https://kraken.iac.rm.cnr.it/C-IMMSIM/) describes the different stages of the process of recognition and response of the immune system to pathogens, while the server simulates three different parts representing three separate anatomical regions in mammals, including the bone marrow, thymus and tertiary lymphoid organs (lymph nodes). The three different intervals are 1, 84 and 168^[35,36]^. In the Xinjiang population, HLA-A*1101, HLA-A*0201, HLA-B*5101, HLA-B*3501, HLA-DRB1*0701 and HLA-DRB1*1501 high frequency alleles were selected for analysis ^[27]^. Finally, the parameters were set to the default values of the software, the simulation parameter random seed was set to 12345, the simulation volume was set to 50 and the simulation step was set to 1050.

#### 1.2.13 Optimization of MEV codons and in silico cloning

We used the online codon optimisation tool ExpOptimizer (https://www.novopro.cn/tools/codon-optimization.html) to analyse and optimise the codons of MEV.For analysis and optimisation, we selected E. coli as the expression host and excluded two restriction endonuclease sites, XHOI and BamHI, and then obtained the DNA sequence of the MEV codon from the results.In silico cloning,we selected pET-28a(+) as the vector. Primers were then designed and polymerase chain reaction(PCR) completed in SnapGene 6.1.1 with primer lengths between 15-30bp, Tm values chosen at 60°C, annealing temperature of 1°C and GC content between 40% and 60%. Finally, the appropriate nucleic acid endonuclease sites (XHOI and BamHI) in the polyclonal site (MCS) region were analysed and the MEV-amplified target gene sequence was inserted into the vector to complete the in-silico cloning.

#### 1.2.14 agarose gel electrophoresis of MEV

Agarose gel electrophoresis of the target gene (after PCR), vector and recombinant plasmid was simulated in SnapGene 6.1.1.and the experiments were completed in TBE buffer and at a concentration of 1% agarose.

#### 1.2.15 Molecular dynamics simulation

iMODS (http://imods.Chaconlab.org/) is the internal coordinate normal mode analysis(NMA) server.To explore the colletive motions of proteins and nucleic acids using NMA in internal coordinates (torsional space)^[37]^.We therefore performed molecular dynamics simulations of the MEV-TLR4 complex and analysed the deformability, stiffness and stability of the complex with the output results.

## Results

### 2.1 Target protein selection

By bioinformatics analysis, the results of antigenicity analysis for VirB8 a nd VirB10 were 0.6221 and 0.6906 respectively both greater than the threshold value of 0.4 with good antigenicity. The combined (Table 1) results show that VirB8 and VirB10 are hydrophilic and stable proteins.

**Table 1.** Basic structure and physicochemical properties of amino acids ino acid.

| amino acid | Number of amino acids | Molecular weight | Instability index | Grand average of hydropathicity(GRAVY) | Theoretical PI |
| --- | --- | --- | --- | --- | --- |
| VirB8 | 239 | 26445.83 | 28.56 | -0.392 | 8.60 |
| VirB10 | 380 | 40442.32 | 36.52 | -0.155 | 7.72 |

### 2.2 Sequence Search

The high precision of the MAFFT in Jalview (Fig.1) suggests that the two proteins share several homologies. Among them, the high homology sequences of VirB8 and VirB10 suggest that these proteins may be derived from the same gene and may play similar roles in the immune response.

**Fig 1.**
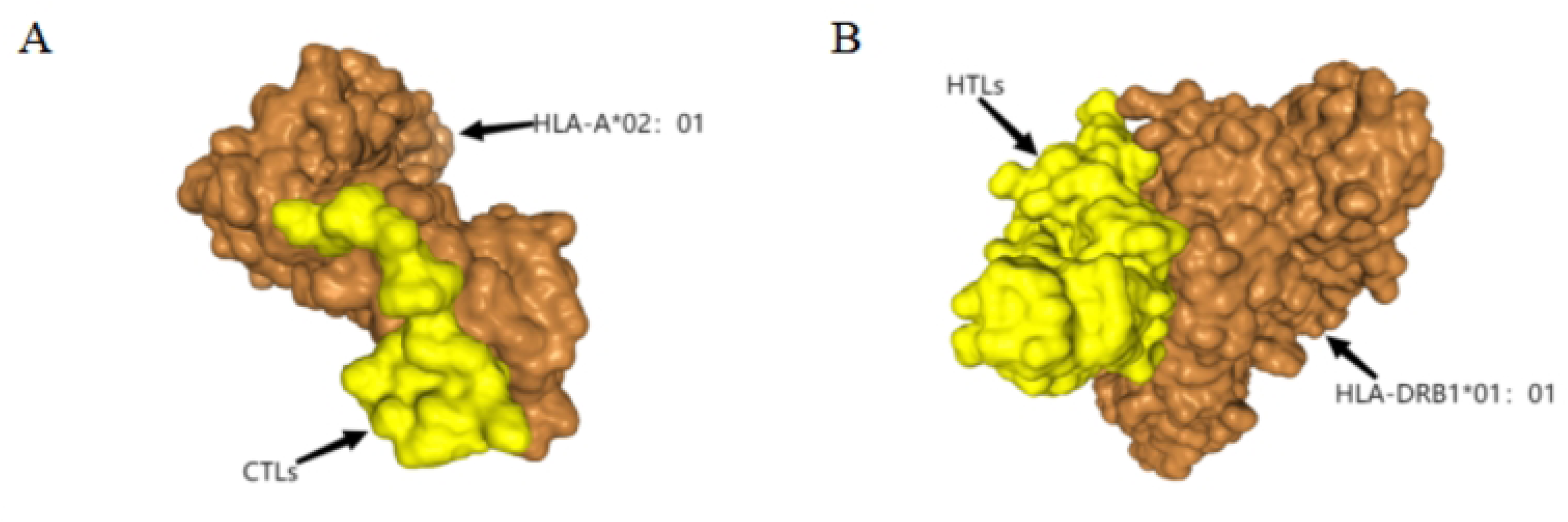
Comparison of protein homologous sequences, the black regions are highly conserved amino acid regions, while the blue regions are similar amino acid sequence regions (darker colours represent high homology)

### 2.3 Prediction of signal peptides

We applied SignalP5.0 and LiPOP1.0 to predict the signal peptide and the final result was no signal peptide for both VirB8 and VirB10 (Fig.2).

**Fig 2.**
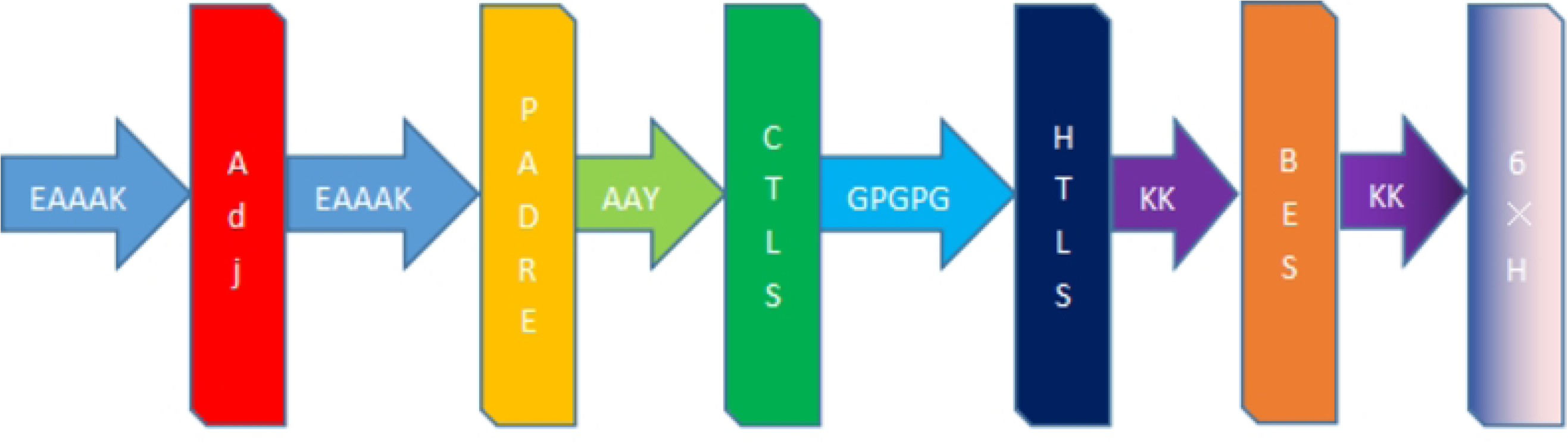
Results of analysis using SignalP-5.0.SP(Sec/SPI) / LIPO(Sec/SPII) / TAT(Tat/SPI) (depending on what type of signal peptide is predicted); CS (t he cleavage site) and OTHER (the probability that the sequence does not h ave any kind of signal peptide) (A) Signal peptide prediction of VirB8: None.(B)Signal peptide prediction of V irB10: None.

### 2.4 Prediction of T-cell epitopes

We selected the top 10 epitopes for each software score and applied VaxiJen to analyse the antigenicity of the epitopes; used AllergenFP v1.0 (http://www.ddg-pharmfac.net/AllergenFP/) to detect whether the epitopes were allergenic; applied ToxinPred (https://webs.iiitd.edu.in/raghava/toxinpred/design.php) to predict the toxicity of the epitopes. Ultimately, two CTL-dominant epitopes and nine HTL-dominant epitopes were obtained (Table 2). These epitopes were highly antigenic, non-toxic and non-sensitizing.

**Table 2.**
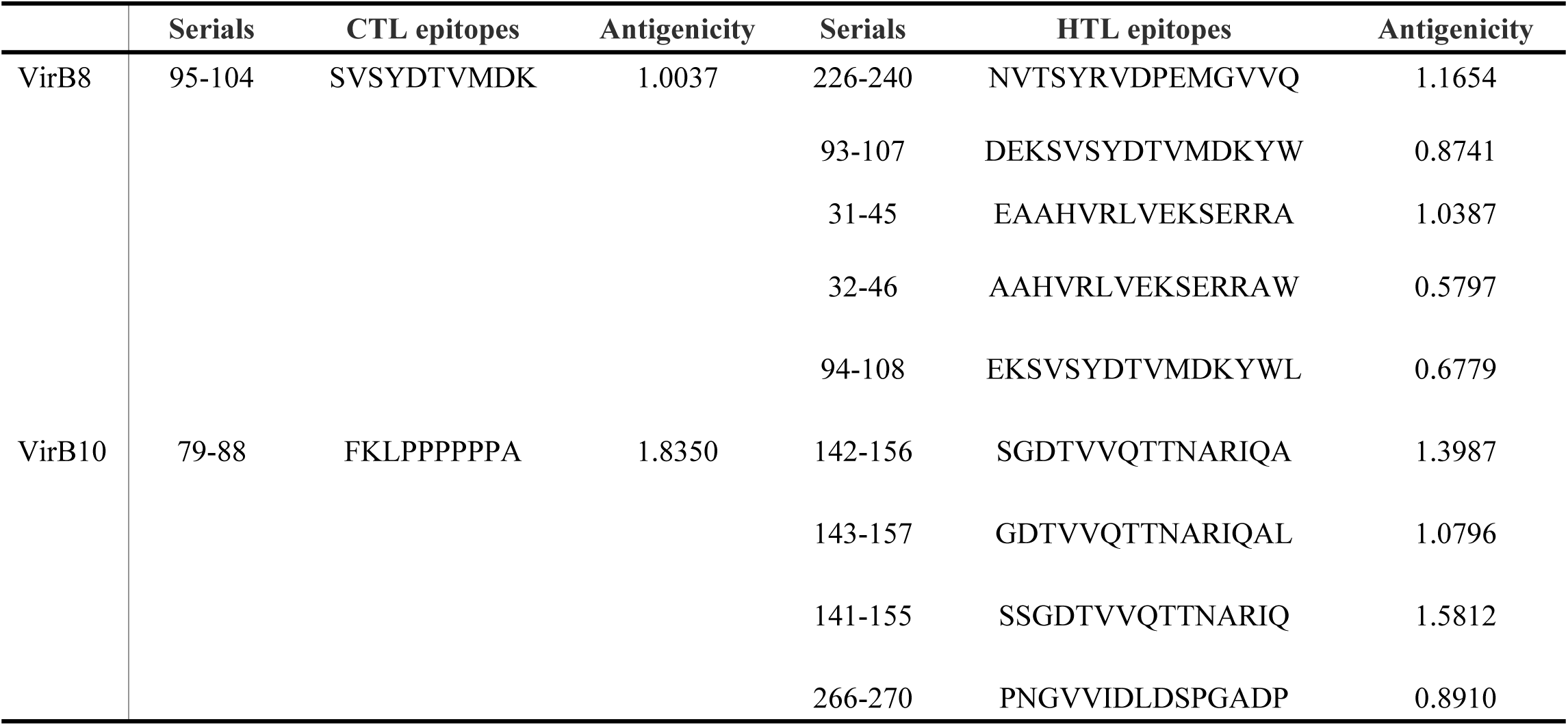
Ultimately selected T cell epitopes(CTL, HTL)

### 2.5 Prediction of B-cell epitopes

By bioinformatics analysis, we obtained six dominant linear B-cell epitopes and six conformational B-cell epitopes (Table 3). Finally, we selected non-toxic and non-sensitizing B-cell conformational epitopes (Fig.3).

**Fig 3.**
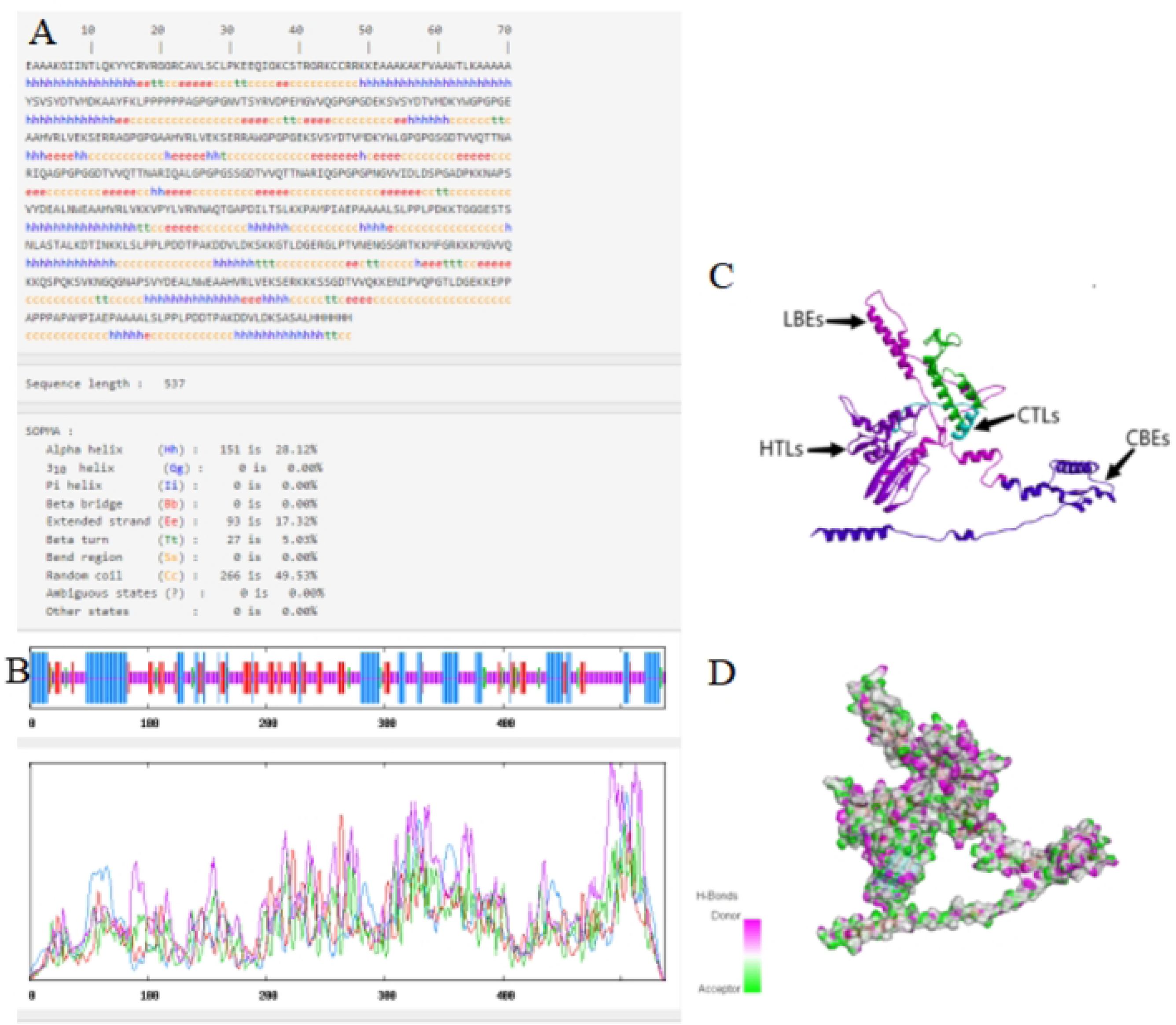
B-cell conformational epitopes. (A-C) B-cell conformational epitope residues of VirB8. (D-F) B-cell conformati onal epitope residues of VirB10 (A)Residues:M1,F2,G3,R4,K5. (B) Residues:M2 35,G236,V237,V238,Q239.(C)Residues:Q6,S7,P8,Q9,K10,S11,V12,K13,N14,G15,Q 16,G17,N18,A19,P20,S21,V22,Y23,D24,E25,A26,L27,N28,W29,E30,A31,A32,H33, V34,R35,L36,V37,E38,K39,S40,E41,R43.VirB10. (D)Residues: K139,S140,S141,G 142,D143,T144,V145,V146,Q147. (E)Residues:E4,N5,I6,P7,V8,Q9,P10,G11,T12,L1 3,D14,G15,E16. (F)Residues: E91,P92,P93,A94,P95,P96,P97,A98,P99,A100,M101, P102,I103,A104,E105,P106,A107,A108,A109,A110,L111,S112,L113,P114,P115,L1 16,P117,D118,D119,T120,P121,A122,K123,D124,D125,V126,L127,D128,S130,A13 1,S132,A133,L134.

**TABLE 3.**
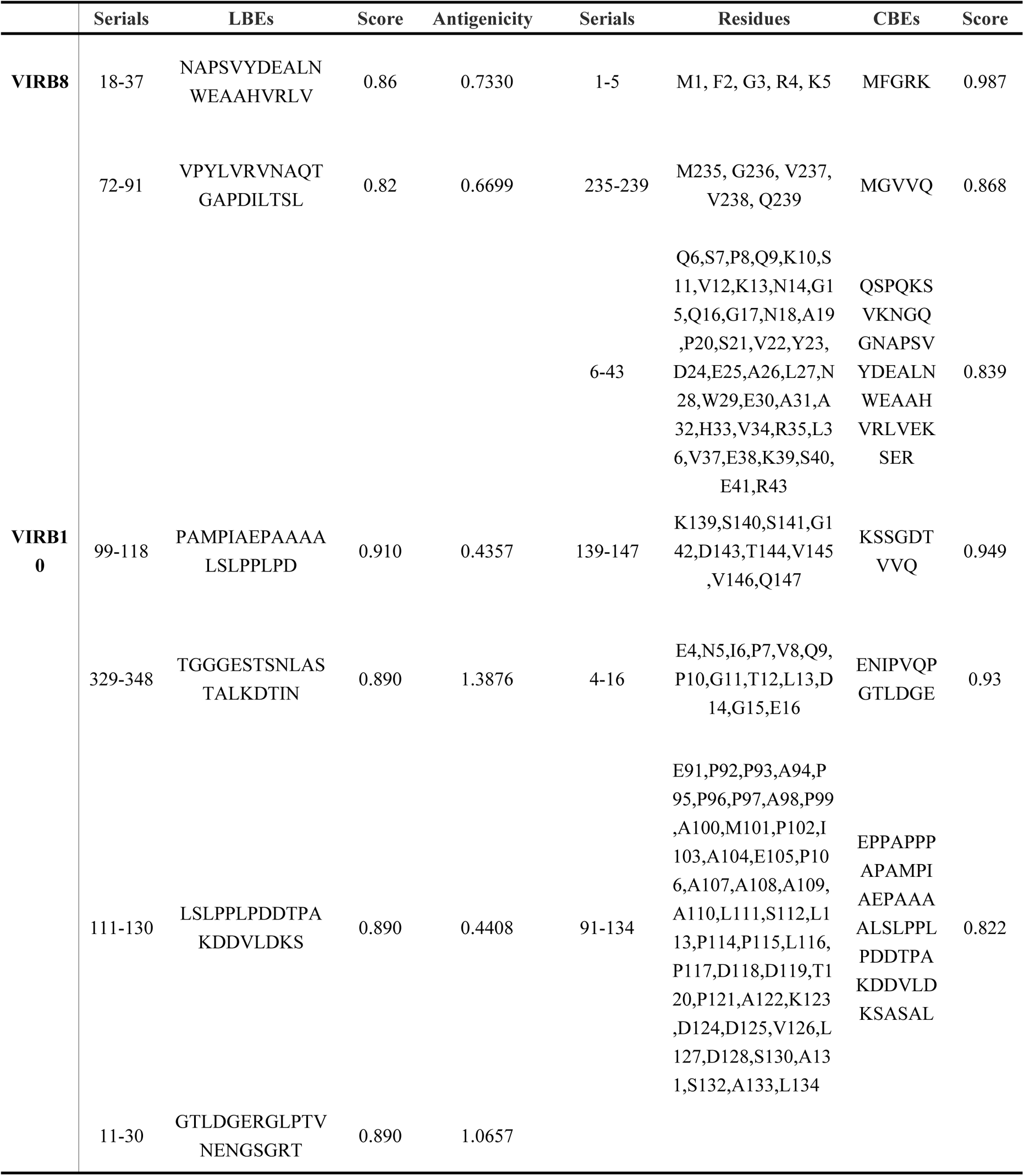
Final selected B cell epitopes(LBES, CBES)

### 2.6 Molecular docking of T-cell epitopes to HLA alleles

Assessing the structural association of HLA alleles with T cell epitopes. Results for CTL epitopes interacting with HLA-A*02:01, Docking Score: –242.47 Confidence Score: 0.8641 Ligand RMSD (Å): 32.94.HTL epitopes interacting with HLA-DRB1*01:01 Results, Docking Score: –275.57 Confidence Score: 0.9249 Ligand RMSD (Å): 150.52. The final indication is that the molecules have good affinity for the docked complexes. (Fig.4).

**Fig 4.**
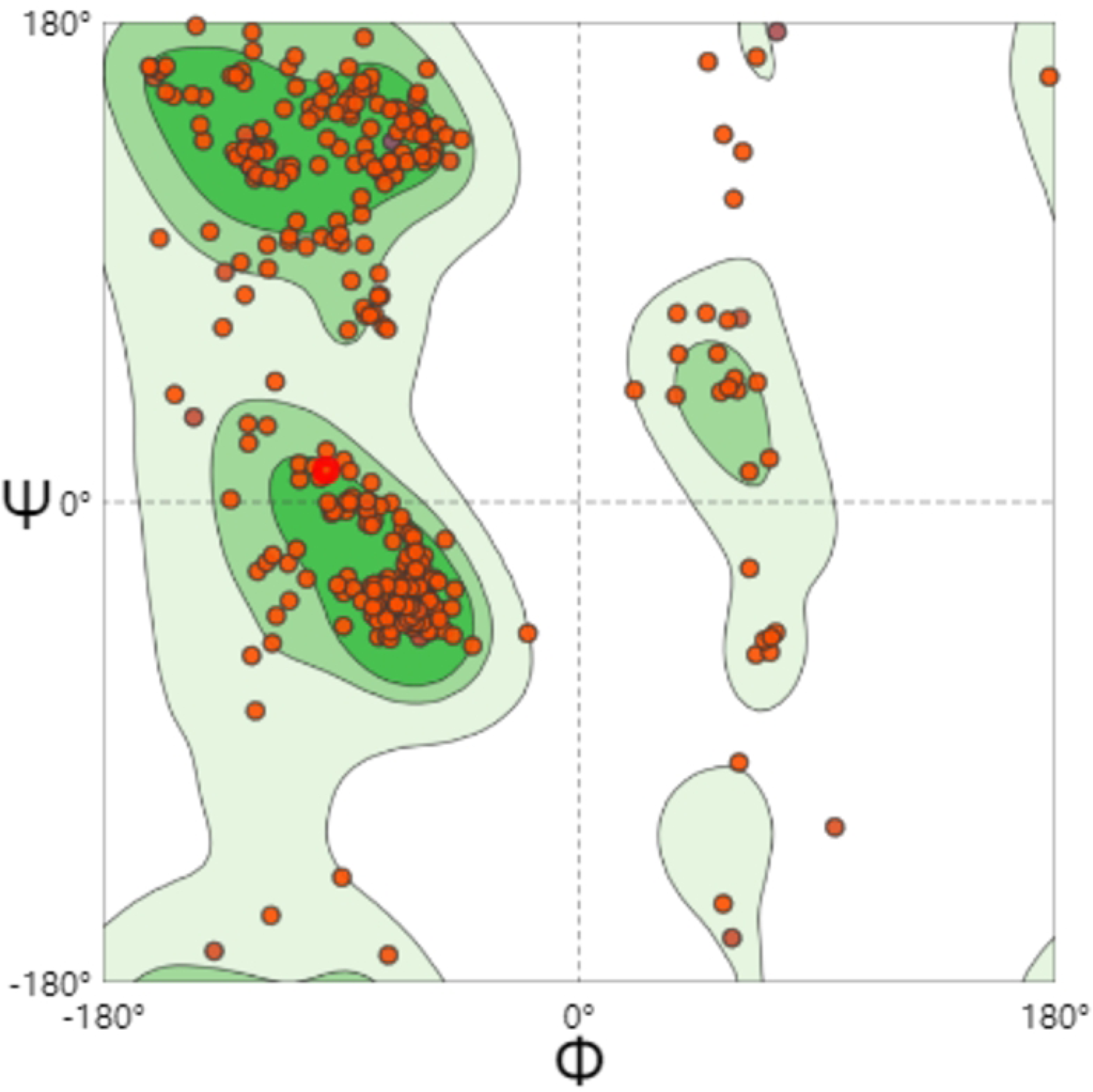
The docked complexes. (A-B) HLA-bacterial peptide complexes. (A) Results of molecular docking of CTL epitopes to HLA-A*02:01. (B) Results of molecular docking of HTL epitopes to HLA-DRB1*01:01.

### 2.7 The construction of MEV

Results of the MEV construction (Fig.5). the dominant epitopes selected for our MEV vaccine construction were 2 CTL epitopes, 9 HTL epitopes, 6 LBE epitopes and 6 CBE epitopes.

**Fig 5.**
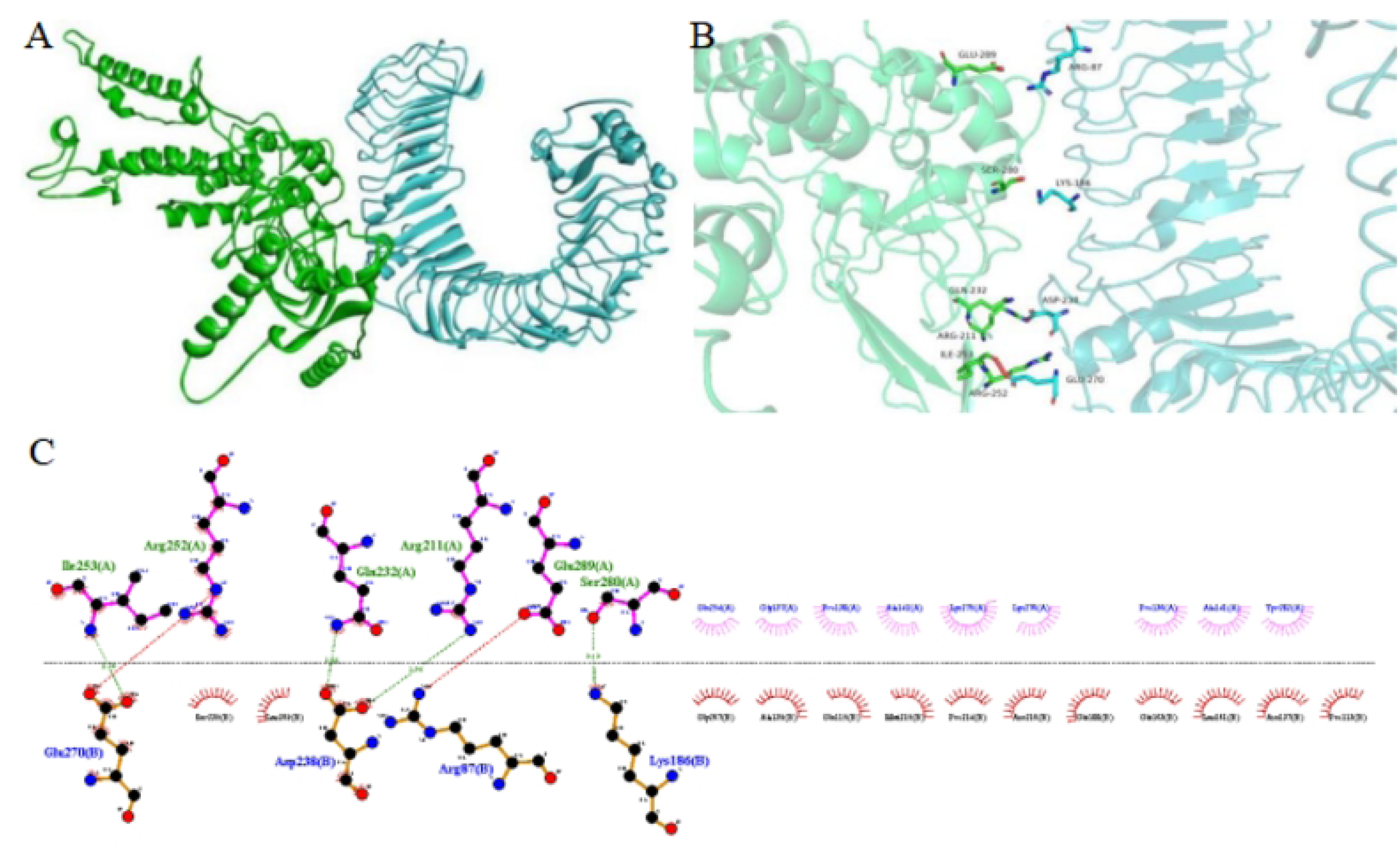
Amino acid sequence of MEV. EAAK, AAY, GPPGPG and KK are linkers.”Adj” in red is β-defensin-3. The Yellow “PADRE” is the PADRE sequence. Dark green”CTLS”represents the dominant epitope of the selected cytotoxic T cells. The dark blue “HTLS” represents the dominant epitope of the selected helper T cells. “BES” represents the dominant epitope of the selected linear and conformational B cells. The “6×H” indicates a polyhistidine tag.

### 2.8 Physicochemical properties, antigenicity, solubility and sensitization of MEV

The molecular formula of MEV was C2479H3998N720O762S14.The molec ular weight of MEV was 56530.22KD. The number of amino acid residues in MEV was 537.MEV had a solubility of 0.99, which means that the protein antigen was soluble.The instability index (II) was computed to be 30.75,less than the threshold 40,so MEV was a stable protein.The GRAVY was –0.626.Indicate s that MEV was a hydrophilic protein.In addition, the antigenicity of MEV is 0.8788 (greater than the threshold value of 0.4), indicating that the protein is a ntigenic. In the sensitisation prediction results it was shown that PROBABLE NON-ALLERGEN.In conclusion, the MEV design is feasible.

### 2.9 Forecasts for secondary and tertiary structures

The predicted results show that in the secondary structure prediction, α-helix accounts for 28.12%, β-turn for 5.03%, random coil for 49.53% and extended strand for 17.32%, the ratio of the four is consistent with the tertiary structure (Fig. 6A, B).Furthermore,we depicted the tertiary structure of MEV in Discover Studio, further demonstrating that the prediction of a tertiary structure is reasonable(Fig. 6C). Ultimately, (Fig. 6D) shows the regions where the donor and acceptor are likely to be present.

**Fig 6.**
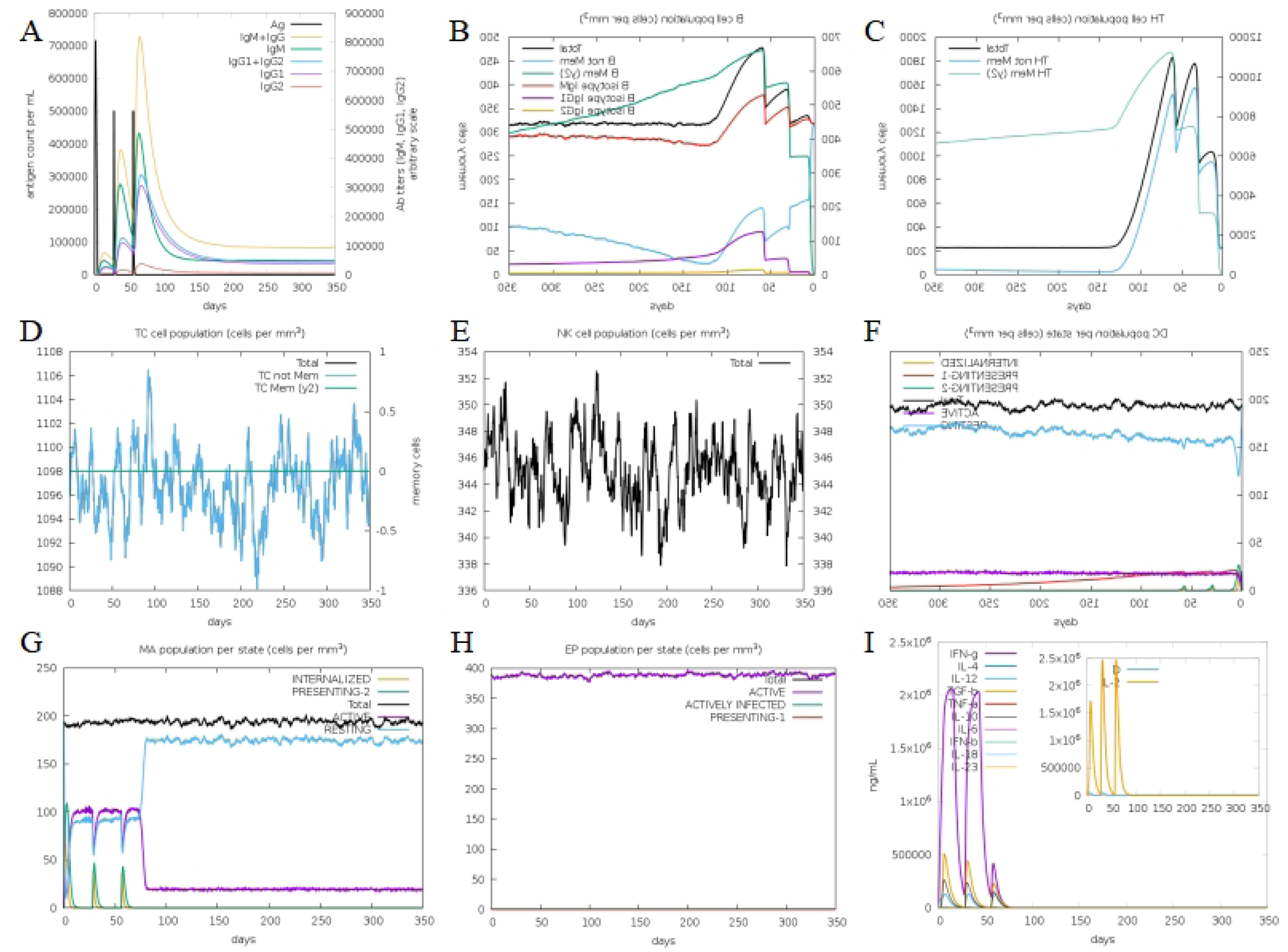
The predicted results of MEV. (A-B) Predicted results for MEV secondary structure. (C)Predicted results for the MEV tertiary structure. (D)H-Bonds of MEV. As illustrated in the figure, the “pink area” stands for the donor, and the “green area” stands for the acceptor.

### 2.10 Quality assessment of models

A Ramachandran plot is a way to visualize energetically favoured regions for backbone dihedral angles against of amino acid residues in protein structure.The number of observed Φ (Phi; C-N-CA-C) / Ψ (Psi; N-CA-C-N) pairs determines the contour lines. (Fig. 7). The dark green region in the Ramachandran plots indicated the allowed region, the light green region in the diagram indicated the maximum allowed region, and the blank region in the diagram indicated the disallowed region.Overall, the quality of MEV’s models was assessed as better.

**Fig 7.**
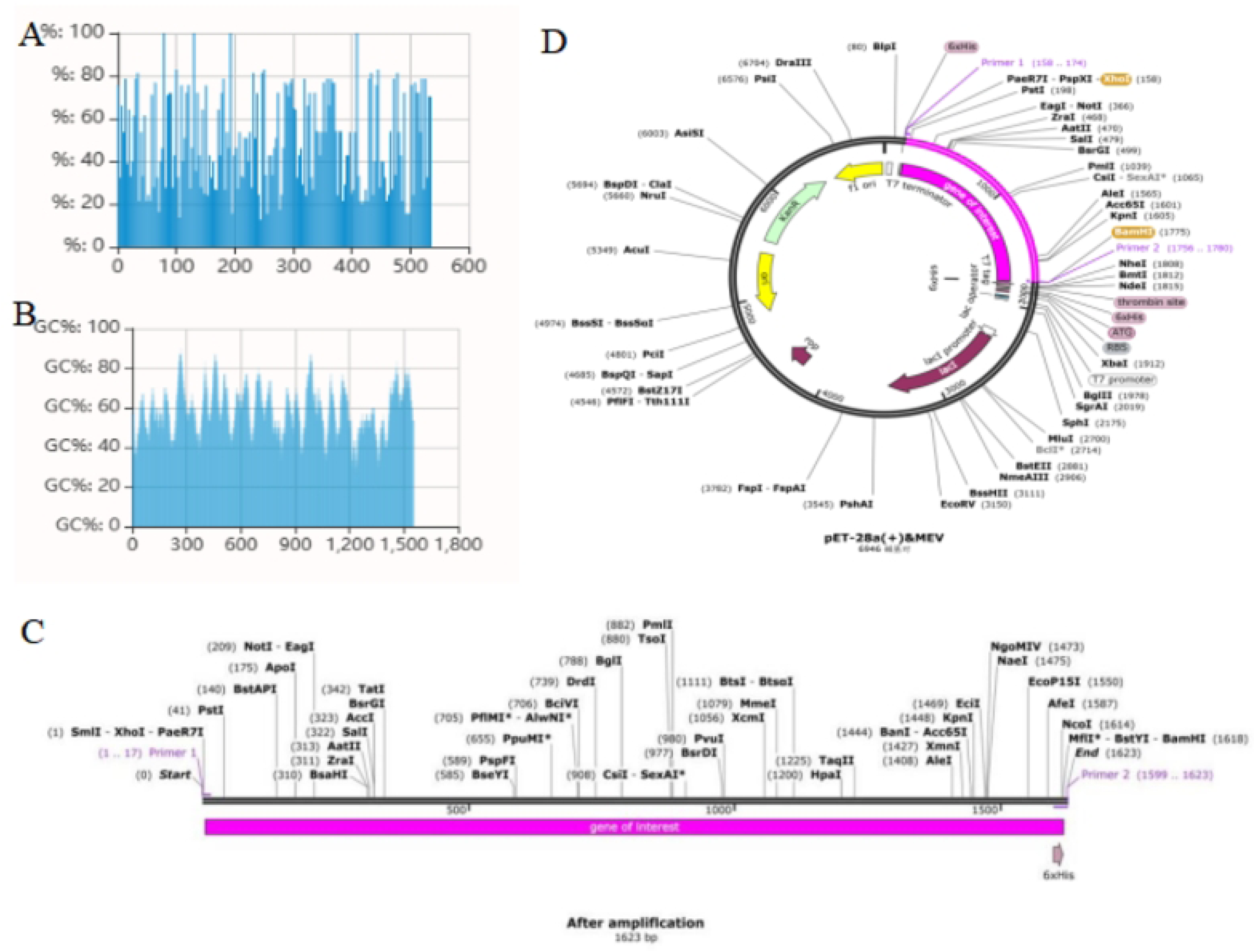
Validation: Ramachandran plot analysis showing 90.47% in favored, 7.10% in allowed, and 2.43% in disallowed regions of protein residues.

### 2.11 Molecular docking

With the HDOCK server we obtained the results of molecular docking. We selected the first place cluster for the analysis of the interaction between receptor (TLR4) and ligand(MEV).The results showed a Docking Score of –349.70, a Ligand RMSD of 54.64 Å and a Confidence Score of 0.9819.Finally, the best docking structure was demonstrated using Discovery Studio (Fig.8A). The 3D interaction structure was then visualised using PyMOL (Fig.8B) and the 2D interaction map was visualised using LigPlot + (Fig.8C).

**Fig 8.**
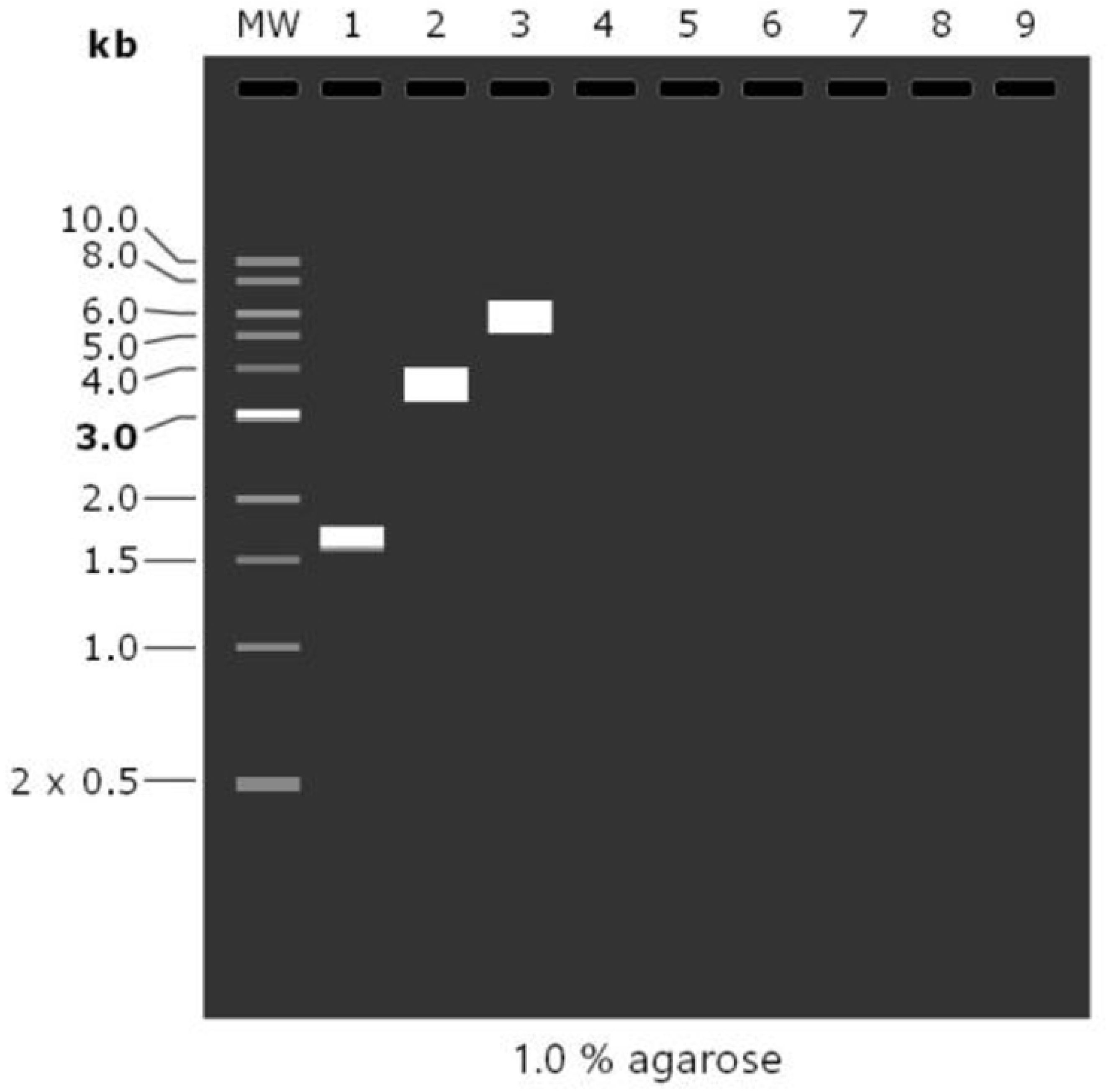
The results of molecular docking. (A)Structural presentation of the MEV-TLR4 complex using Discovery Studio: MEV donor in green and TLR4 acceptor in blue. (B) Analysis of the interaction of the MEV-TLR4 complex and its 3D image taken using PyMol. (C) Analysis of the interaction of the MEV-TLR4 complex and its 2D image taken using Ligplot.

### 2.12 Immunosimulation

The C-ImmSim server was used to simulate the immune response to three injections of the vaccine.In the secondary and tertiary immune response, the concentrations of IgM and IgG continued to rise as the antigen decreased, and the amount of IgM was consistently higher than IgG, peaking at the tertiary response (Fig.9A).B cells are mainly involved in humoral immunity and play an important role in the stimulated immune response, with the number of B cells increasing with the three doses of vaccine and eventually reaching a peak (Fig. 9B).The growth trend of helper T cells after three doses of vaccine was similar to that of B cells, eventually reaching a peak (Fig. 9C).Macrophage activity was enhanced with the three doses of vaccine (Fig. 9G).In contrast, cytotoxic T cells (Fig. 9D), natural killer cells (Fig. 9E), dendritic cells (Fig. 9F) and EP (Fig. 9H) all showed relative stability in general.In addition, the vaccine injection induced a high response of cytokines and interleukins, resulting in significant elevations of IFN-γ, TGF-β, IL-10, IL-12 and IL-2. finally,the danger signal was extremely low, indicating a difference in immune response. (Fig. 9I)

**Fig 9.**
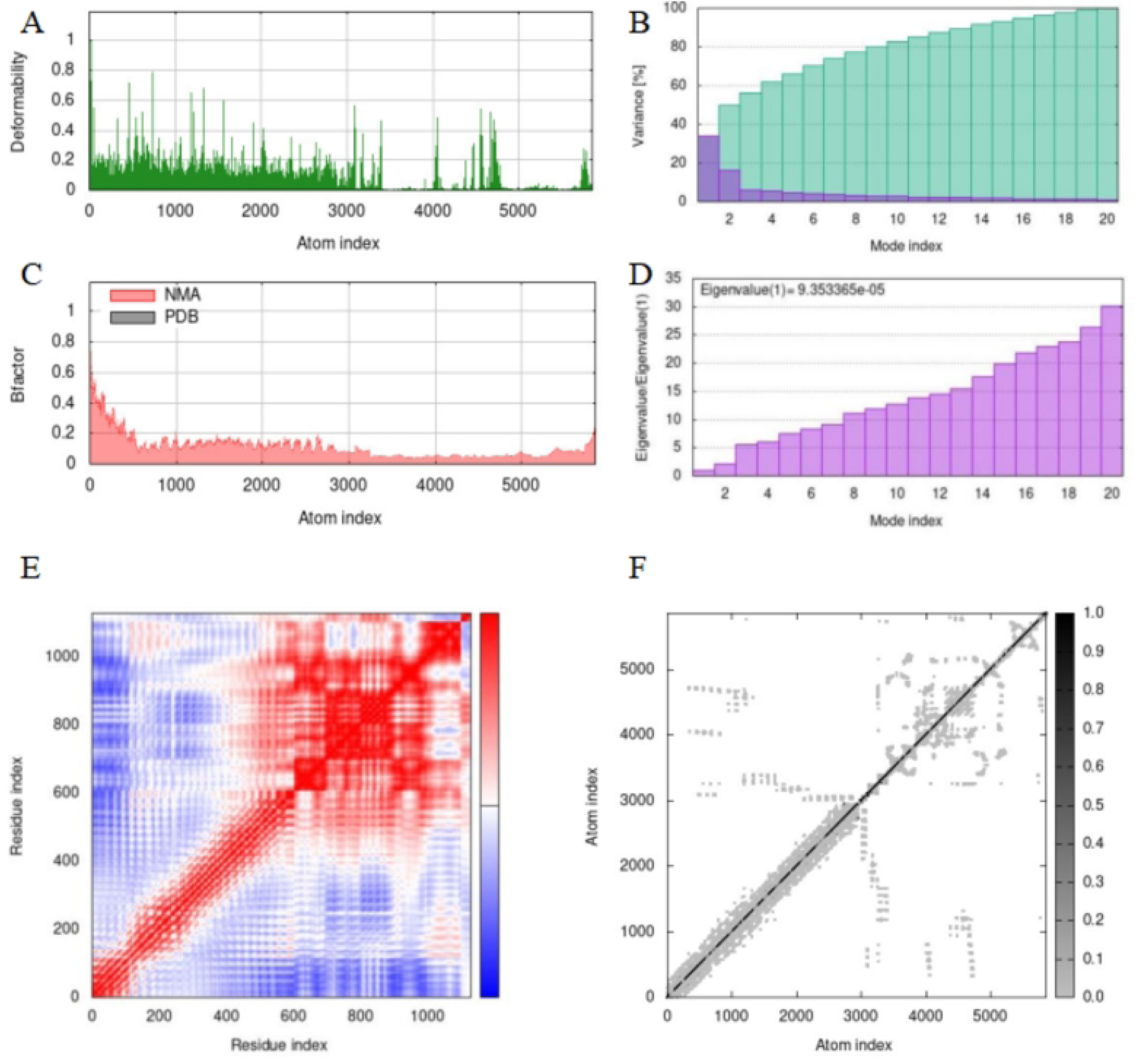
The results of C-ImmSim. (A) The immunoglobulin production after antigen injection.(B) The B cell population after three injections.(C)The Helper T Cell Population after three injections.(D) The Cytotoxic T Cell Population after three injections.(E)The NK-cell Population after three injections.(F)The Dendritic cell Population per state after three injections.(G)Macrophage Population per state after three injections.(H)The EP Population per state after three injections.(I)Concentration of cytokines and interleukins. Inset plot shows danger signal together with leukocyte growth factor IL-2.

### 2.13 Optimization of MEV codons and in silico cloning

The quality of the codon optimisation was measured by the display of the codon adaptation index (CAI) and GC content. the optimized CAI value for MEV was 0.80. the ideal range for GC content was 30%-70% and the optimized GC content for MEV was 58.35%, which was within the ideal range (Fig.10A, B). Based on the principles of primer design, a forward primer (5′-CTCGAGGAAGCGGCGGC-3′) with a length of 17, a Tm value of 62 and a GC content of 76%; and a reverse primer (5′-GGATCCATGGTGGTGATGATGGTGC-3′) with a length of 25, a Tm value of 63 and a GC content of 56% were designed. The target gene for MEV was then amplified in SnapGene(Fig.10C). Restriction endonuclease sites XHOI and BamHI were inserted into the N and C ends of the optimised codons at the time the primers were designed. Finally, these codons are inserted into the MCS structural domain in the vector and should be considered to correspond to the restriction endonuclease sites (XHOI and BamHI) on the vector (Fig.10D).

**Fig 10.**
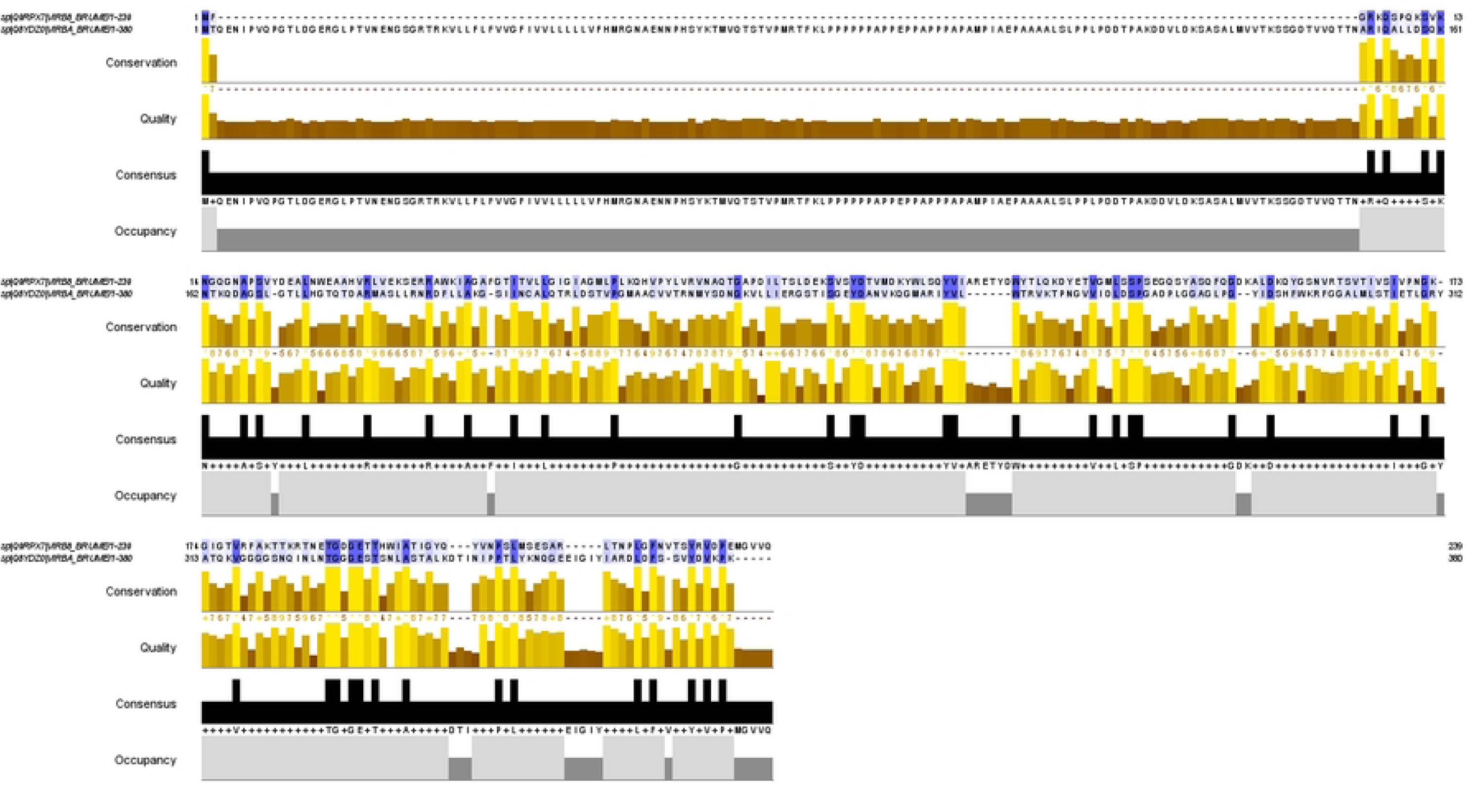
Codon optimization of MEV and construction of plasmid vectors. (A) CAI after codon optimization: 0.80. (B) GC content after codon optimization: 58.35%. (C) MEV after polymerase chain reaction. (D) The pink sequence (gene of interest) is the MEV codon sequence optimized after insertion into the vector (pET28a (+)). The cloning was done in SnapGene6.1.1.

### 2.14 agarose gel electrophoresis of MEV

As shown in the figure (Fig.11). The amount of each DNA was consistent with previous predictions, with 1611bp of MEV sequence,with 1623 bp of MEV sequence after PCR, 5639 bp of pET-28a(+) sequence, and 6946 bp of recombinant plasmid sequence.

**Fig 11.**
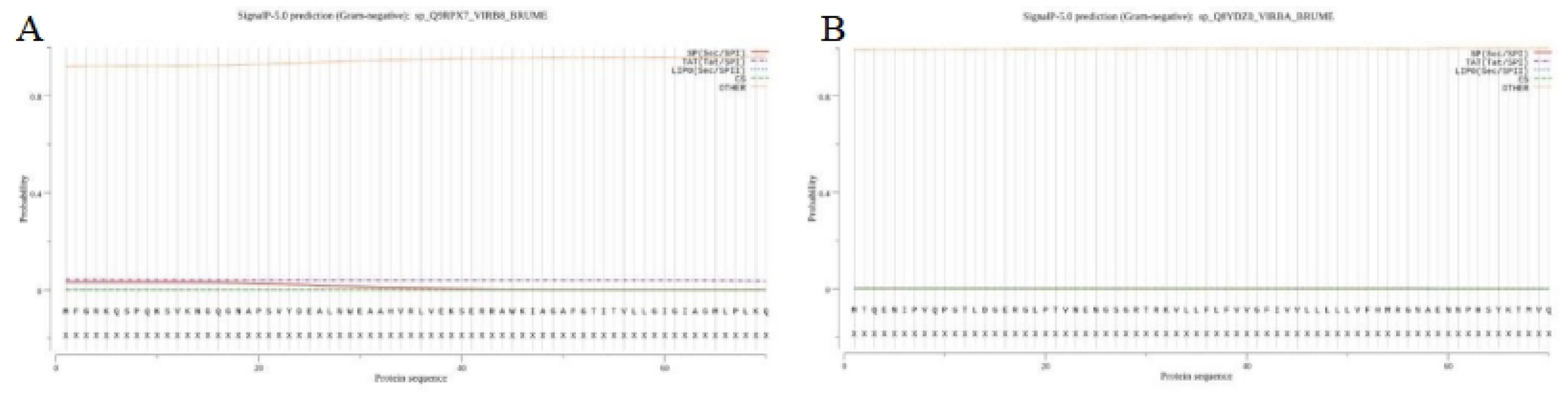
Mock agarose gel electrophoresis results. “1” represents MEV-PCR, “2” represents pET-28a (+), “3” represents pET-28a (+) &MEV recombinant plasmid.

### 2.15 Molecular dynamics simulation

The molecular motion was analysed by normal modal analysis using IMODs and the results are shown in (Figures 12). The main-chain deformability is a measure of the capability of a given molecule to deform at each of its residues. The location of the chain ‘hinges’ can be derived from high deformability regions (Fig.12A). The variance associated with each normal mode is inversely related to the eigenvalues. Coloured bars show the individual (purple) and cumulative (green) variances. The variance plots show the progressive decline of individual variances (Fig.12B). The experimental B-factor is taken from the corresponding PDB field and the calculated from NMA is obtained by multiplying the NMA mobility by (8pi^2). The relationship between the NMA and PDB regions in the complex is depicted in the B-factor diagram(Fig.12C).The eigenvalue associated to each normal mode represents the motion stiffness. Its value is directly related to the energy required to deform the structure.The eigenvalue in the eigenvalue plot is 9.353365e-05, which, according to previous studies, indicates that the complex has low deformability and good stability(Fig.12D).The covariance matrix represents the coupling between pairs of residues and the way in which certain parts of the macromolecule move, i.e. whether they experience correlated (red), uncorrelated (white) or anti-correlated (blue) motions(Fig.12E).The elastic network model defines which pairs of atoms are connected by springs.Each dot in the graph represents one spring between the corresponding pair of atoms.Dots are colored according to their stiffness.A darker shade of grey indicates higher stiffness, i.e. each chain of the MEV-TLR4 complex has a higher stiffness(Fig.12F), which indicates a more stable complex.

**Fig 12.**
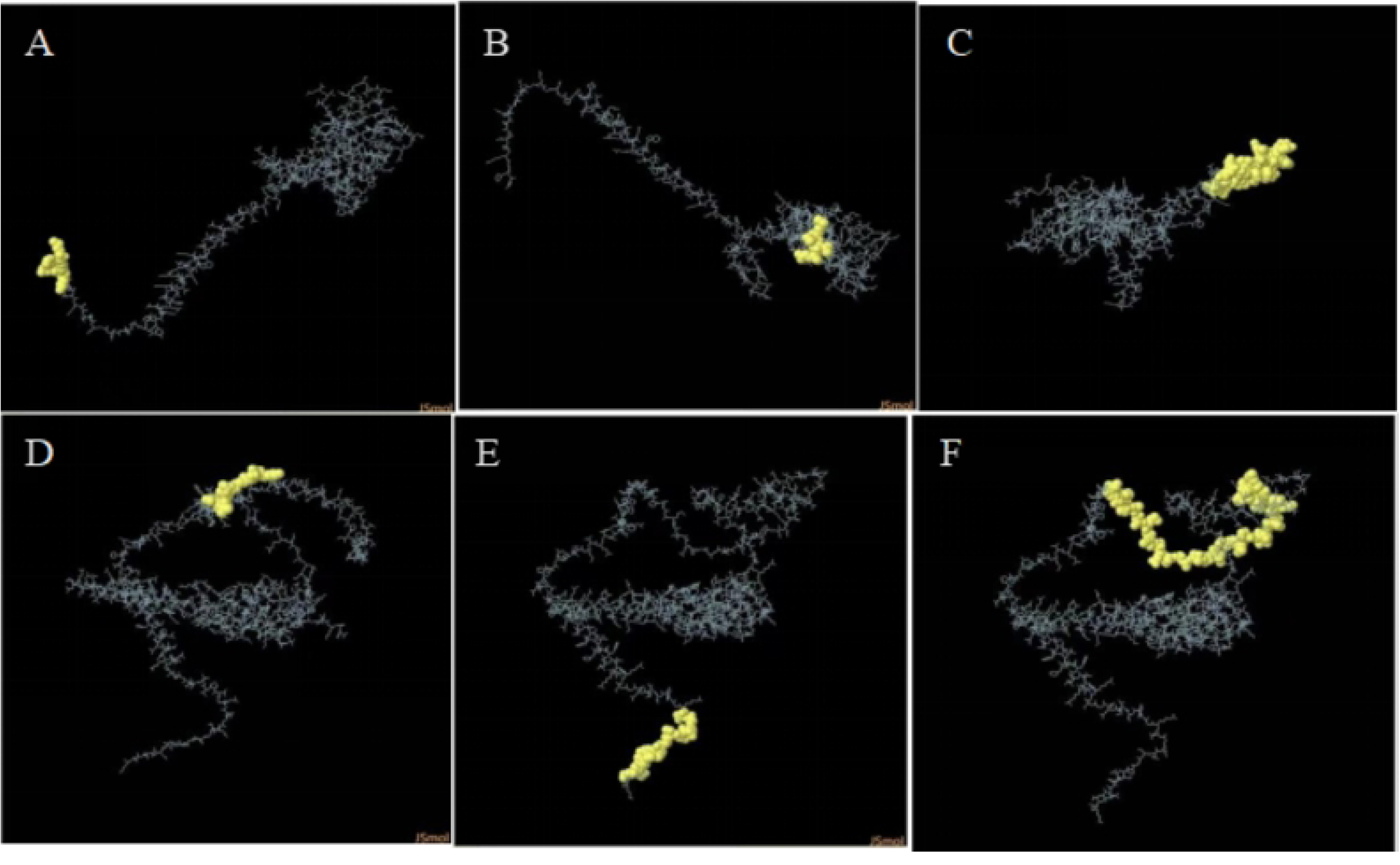
The results of IMODs. (A-F) Molecular dynamics simulation results. (A) Deformability values. (B) The variance associated to the modes. (C) B-factor and NMA graph. (D) Eigenvalues plot. (E) Covariance matrix graph. (F) The elastic network models.

## Discussion

Brucellosis is a debilitating zoonotic disease that can cause significant economic losses in livestock populations worldwide^[38,39]^.As there is currently no brucella vaccine for humans, it is vital to develop a more effective and safer vaccine for humans^[40]^.The type IV secretion system (T4SS), encoded by the virB manipulator, is an important virulence factor for Brucella abortus, while its core component consists of VirB6-VirB1016.VirB8 is a two-site endosomal protein that plays a critical role in the nucleation of T4SS channels^[41,42].^VirB10 is a bilayer protein inserted into the bacterial endosome, and the proline-rich region plays a key role in core complex assembly and substrate secretion^[43,44]^.

Following previous studies, the proteins required to construct novel MEVs must be highly antigenic ^[45]^.In addition, stability, hydrophilicity and allergenicity have been assessed.VirB8 and VirB10, the core components of the type IV secretion system (T4SS), were selected to construct a novel multi-epitope vaccine. The homology of the two proteins was verified during sequence alignment and met the requirements for novel vaccine design.There are no reports of MEV construction based on two proteins, VirB8 and VirB10.

Signal peptides usually contain 15-30 amino acids^[46,47]^They are usually located at the N-terminal end of the protein and influence the start of protein translation, and the different primary structures of the signal peptide even influence protein folding and translocation^[48,47]^,The expression level of a protein can be altered by replacing the signal peptide^[49,47]^.In our study, signal peptides were predicted for VirB8 and VirB10 using SignalP5.0 and LiPOP1.0.The results showed that neither protein had a signal peptide and we would not need to remove the signal peptide sequence deliberately. Therefore, we believe that we can proceed to the next step of the analysis.

The main goal to be achieved with vaccines is to provide lasting memory. It is therefore crucial to activate B cells and T cells to achieve this aspect.To predict the epitopes of CTL, HTL, LBE and CBE and to select suitable candidate vaccines, different databases and online servers were used^[50,51]^,Helper T lymphocytes initiate humoral and cell-mediated immune responses, cytotoxic T lymphocytes prevent virus transmission by killing virus-infected cells and producing antiviral cytokines, and B lymphocytes are primarily involved in humoral immune responses^[52,53]^.

A multi-epitope vaccine consisting of CTL, HTL and B-cell epitopes triggers broad immune protection, i.e. induces activation of cellular and humoral immune responses and enhances their immunogenicity^[54,55,56]^.We obtained 2 dominant CTL epitopes from both proteins using IEDB and NetCTLpan1.1 server, 9 dominant HTL epitopes using IEDB and NetMHC-IIpan-4.0, 6 B-cell linear epitopes using ABCpred and 6 B-cell conformational epitopes using Ellipro of IEDB. cell conformational epitopes using Ellipro of IEDB.Our MEV was then constructed by selecting the superior epitopes obtained above.In the vaccine construction, the dominant epitopes are connected by linkers.We linked the CTL, B-cell and HTL epitopes to the AAY, KK and GPGPG linkers, respectively.The linker ensures that each epitope can trigger the immune response independently and avoids the creation of new epitopes that interfere with the immune response induced by the original epitope^[57]^.However, the immunogenicity of multi-epitope vaccines is poor when used alone and requires adjuvants for coupling^[58]^.Adjuvants are important components of vaccine formulations, preventing infection and influencing the specific immune response to antigens, maintaining the stability of peptides and enhancing their immunogenicity^[59]^.To improve the immunogenicity of MEV, the adjuvant human beta-defensin-3 (hBD3) was fused to the N terminus with the help of the EAAAK linker^[60]^,Immediately followed by access to the PADRE sequence to reduce the role of human HLA-DR polymorphisms^[61]^,Finally, the histidine sequence was added to obtain the complete MEV.Molecular docking between the HLA allele and the T cell epitope demonstrates the good affinity of the docking complex.

In structure-based reverse vaccinology, the protein molecular weight of our designed vaccine is 56 KD, which is in the ideal range (<110 KD)^[62]^.The theoretical pI of the vaccine construct was 9.39 and the number of amino acids was 537, indicating the basic nature of the vaccine construct.Instability index and GRAVY values indicate vaccine protein stability and hydrophobicity.Additionally,assessment of sensitization and antigenicity showed that the vaccine was immunogenic and highly antigenic (antigenicity of 0.8788 < 0.4) and that it was not allergenic.These results show that our vaccine constructs are stable, hydrophilic, antigenic, soluble and non-sensitising.In the next secondary structure predictions, β-turns and random coils account for 5.03% and 49.53% respectively.The high proportion of beta-turned and random coils in MEV suggests that the vaccine protein may form antigenic epitopes^[63]^.The tertiary structure of the MEV was predicted by the RoseTTAFold server and the quality of the tertiary structure of the MEV was verified by the SWISS-MODEL structural assessment service.The results show that the three-stage structure of the MEV has a high degree of accuracy and a high approximation factor, and that the overall structure is reliable and of good quality.However,the predicted β-turn angles and random coils are consistent with the secondary structure predictions, which further suggests that our vaccine constructs are correct.Strong interaction between antigenic molecules (MEV) and immune receptor molecules (TLR4) is necessary to initiate an immune response^[64,65]^.Toll-like receptor 4 (TLR4), an innate immune receptor, is commonly involved in multi-epitope vaccine construction^[66]^,Protein-ligand docking analysis and molecular dynamics simulations were performed on the MEV-TLR4 complex to examine the stability between the protein and TLR-4 and to calculate the potential immune response.In the atomic interaction diagram it is shown that there are strong interactions between molecules so that they can be transported throughout the host body^[67]^.We then used MD simulations to explore the stability of the complexes, generating eigenvalue data showing the stiffness and energy required to move the docked complexes.The results indicate that MEV binds stably to TLR4.To verify that this structure can be involved in the humoral and cellular responses studied, we performed immune simulations of vaccine effects^[68]^.With three vaccinations, we found that T and B cells in the body increased with the number of injections and peaked at the third vaccination.Furthermore, MEV increased the levels of cytokines (IFN-γ,TGF-β, IL-10, IL-12,IL-2), IgG and IgM.IFN-γ indicates cell-mediated immunity, a chemokine that supports B cell proliferation, Ig isotype switching and humoral responses.Antigen-presenting cells display HTL epitopes when using MHC class II molecules, and the HTL epitopes produce associated cytokines (IFN-γ, IL-10) to kill pathogens until they are completely eliminated^[69,70,71]^.These results suggest that MEV can be designed to trigger a robust immune response without producing an allergic reaction and could be considered an excellent candidate for a brucellosis vaccine.

Efficient expression of MEV vaccine protein in the Escherichia coli system is essential for the production of recombinant proteins^[72]^,We use the online codon optimisation tool ExpOptimizer to optimise the amino acid sequence of the vaccine for various parameters such as 5’ region optimisation (translation initiation efficiency), DNA repeat optimisation and GC content optimisation^[73,74]^.The optimised codon GC content (58.35%) and codon adaptation index (CAI=0.80) showed a good probability of vaccine protein expression levels in E. coli hosts.XhoI and BamHI restriction sites were then added to the 5′ and 3′ ends of the codon sequence and primers were designed for them to facilitate the polymerase chain reaction of the target gene.The final vaccine sequence was then cloned into the pET28a(+) vector, yielding a 6946bp recombinant plasmid. Eventually,mock agarose gel electrophoresis experiments were performed on the target gene, vector and recombinant plasmid. Animal experiments need to be refined for this study to follow.

Overall, MEV exhibits desirable physicochemical properties and immune response.Molecular dynamics simulations demonstrate the high stability of MEV. Immunosimulations show that MEV triggers an immune response consistent with our hypothesis.In summary, the novel MEV we constructed can be used as a candidate vaccine for brucellosis and provide a theoretical basis for the development of future brucellosis vaccines.

### DATA AVAILABILITY STATEMENT

The data derived from public domain information: Uniprot database (https://www.uniprot.org/) and PDB library (https://www.rcsb.org/). The data that support the findings of this study are available in the methods and/or supplementary material of this article. The data that support the findings of this study are available from the corresponding author upon request. There are no restrictions on data availability. If you have any questions, please contact me.

## Abbreviations

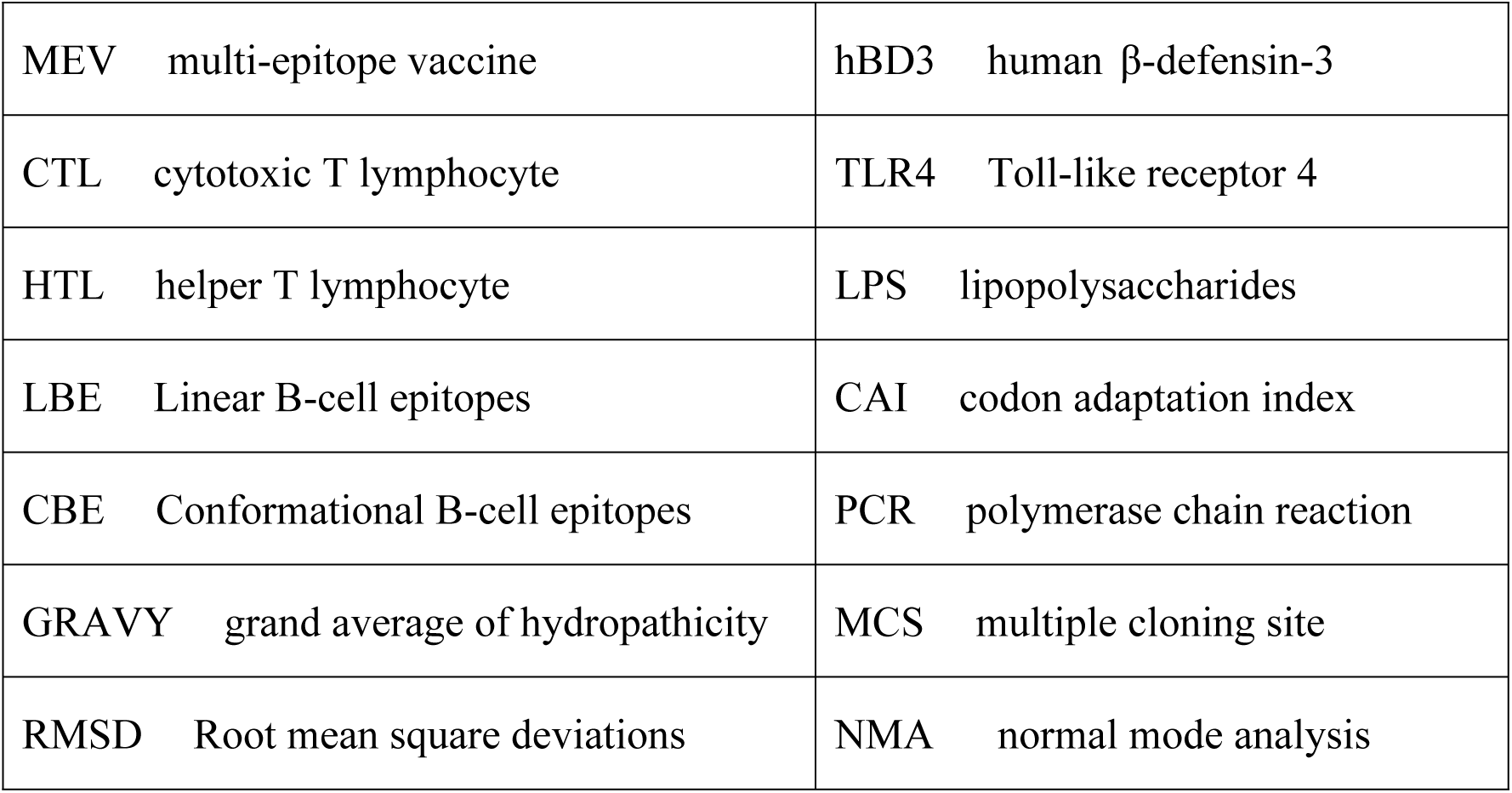

## ACKNOWLEDGMENTS

Te authors are thankful to the State Key Laboratory of Pathogenesis, Prevention, Treatment of Central Asian High Incidence Diseases, Te First Afliated Hospital of Xinjiang Medical University, PR China.

## AUTHORS’ CONTRIBUTIONS

This study was conceived and designed by Jianbing Ding and Fengbo Zhang. Bioinformatic analysis was performed by Zhengwei Yin, Min Li, Ce Niu, Mingkai Yu, Xinru Xie, Gulishati Haimiti, Wenhong Guo, Juan Shi and Yueyue He. The manuscript was drafted by Zhengwei Yin and edited by Jianbing Ding and Fengbo Zhang.All the authors contributed to the article and approved the manuscript.

## ORCID

Jianbing Ding: http://orcid.org/0000-0001-5506-7665

Fengbo Zhang: https://orcid.org/0000-0001-5795-8672

